# Canary Mating Season Songs Move Between Order and Disorder

**DOI:** 10.1101/2024.10.06.616872

**Authors:** S. Levin, Y. Cohen

## Abstract

Many complex behaviors involve sequences of basic motor or vocal elements governed by syntactic rules, which facilitate flexible and adaptive actions. Songbirds that crystallize their repertoire of vocal syllables and transitions allow researchers to build probabilistic models of syntax rules, providing insight into underlying neural mechanisms. However, in many complex behaviors, syntax rules change over time, such as during learning or in response to new environmental and social contexts.

In this study, we investigated the songs of canaries, a seasonal songbird species. Canaries learn a repertoire of 30-50 syllable types, produce syllables in repeat phrases, and organize these phrases into sequences with long-range syntactic dependencies. Since canaries are known to adapt their repertoire annually, we recorded their songs during the spring mating season and examined the syntactic properties that determine syllable sequencing. Over days and weeks, we observed changes in syllable usage rates, syllable numbers within phrases, phrase positions in songs, and in the long-range dependencies of phrase transitions. Acoustic features of syllables were also found to shift alongside these syntactic changes.

Quantifying the variability of these properties revealed that the observed changes were not random. Most birds exhibited a clear trend of moving between order and disorder in their song’s syntactic and acoustic features. Interestingly, this trend varied across individuals; some birds increased their stereotypy and decreased variability across days, while others adopted more disordered and variable song structures.

These findings establish canaries as a valuable animal model for studying the neural mechanisms underlying syntax rules in complex motor sequences, their plasticity in social and environmental adaptation, and in implementing individual-specific strategies.

## 1 Introduction

Many complex behaviors are sequences of basic elements such as motor gestures or vocal syllables. With practice, the individual elements may be very precise [11, 7, 8, 3], even stereotyped, but their sequential order in behaviors ranging from brushing teeth to our highly evolved speech, tool-use, and dance, is often variable across renditions. This sequence variability creates rich repertoires - affording adaptive and flexible skills that are beneficial to the organism [2, 5, 6].

Some sequences are produced often and others not at all in these behavior repertoires. For example, when brushing teeth we may freely choose to brush one side of the mouth before the other but we will never put toothpaste on the brush at the last step. These regularities are captured by sensorimotor syntax rules that govern the probability of element-to-element transitions [1, 26, 10, 48].

Syntax rules reflect key cognitive functions such as working memory [39, 44, 41, 43], motor planning [42, 32, 34, 13], and on-the-fly improvisation. To describe syntax rules that govern transitions in animal behaviors, researchers use mathematical models like Markov models, probabilistic graphs [31], and finite state automata [10, 48] and then apply these models to answer questions about both behavior and its underlying neural mechanisms. For example, transition probabilities in rodent grooming sequences were shown to be encoded by neural activity in motor cortex and dorsomedial striatum neurons that differed between habitual and rare transitions [42]. The outcome probabilities of syllable-to-syllable transition in bird songs were shown to depend on transitions that occurred several seconds prior in the same song [31, 39] or on the outcome of the same ’branching’ transition in the previous song [29].

In these studies, and in studies that extract sequences from continuous motion [33, 50], syntax rules were either assumed to be fixed or were shown to be actively maintained (e.g. in the songs of Bengalese finches [29]). But syntax rules can change: during learning, in response to environmental changes [29, 44], and also to pursue intrinsic goals like a jazz player that changes her improvisation rules trying to reach new audiences. In these changes, transitions between motor elements may change their dependence on sequence context and occur alongside variation in the motor elements themselves [25]. Understanding neural systems that generate, and also change, syntax rules will advance systems neuroscience and potentially computer science, specifically large language models. Still, little has been done to study dynamic syntax rules in animal models.

Here, we study the courtship song of the canary, a seasonal songbird, and find that both canary syntax rules and syllable acoustics vary across days and weeks of their spring mating season. Canary time scales and syntax complexity are similar to those found in human speech: Canary songs are typically 10-40 seconds long (some last more than a minute) and compose of up to 50 different syllable types (Fig. 1A). Some syllable types appear in more than 50% of the songs while others are rare and appear in less than 5% of the songs (Fig. 1B). A typical song contains hundreds of syllables produced in repeats, called phrases, each lasting 1-2 seconds and sung preferentially in the early, middle, or late third of the song (Fig. 1A,C). The number of syllable repeats per phrase varies between songs and the phrase sequences follow long-range syntax rules - sequence dependencies spanning several seconds (Fig.1D-F) [31, 39] and bridging up to 50 syllables of ongoing song. [29, 28, 27].

**Figure 1:**
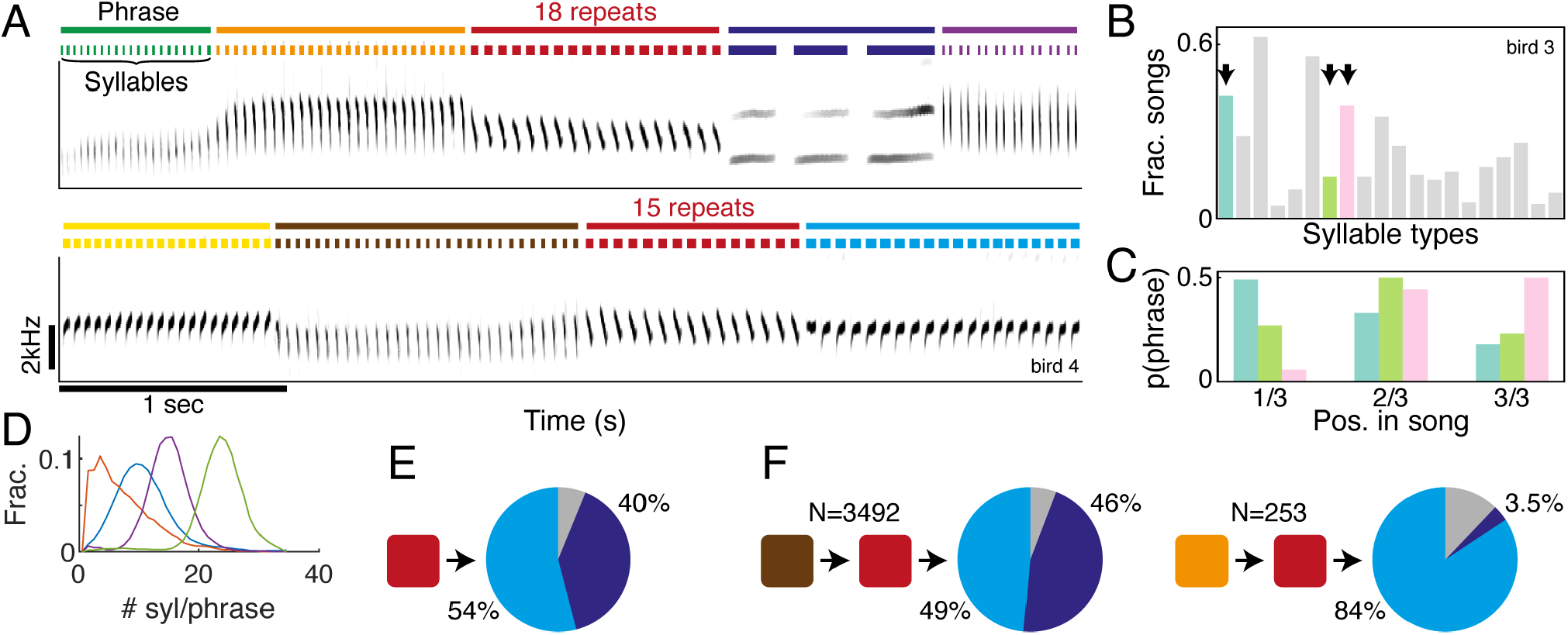
Canary syntax rules are defined by syllables with different probabilities of usage, repeat number, position in song, and transitions. **A.** Sonograms showing two segments from an example canary song (from bird 4). Colored bars annotate the time and identity of syllables (bottom bars) and their repeat phrases (top bars). The dark red phrase appears twice in this song and contains different numbers of syllable repeats. **B**. Bars show the fractions of bird 3’s songs that include different syllable types (x-axis). Arrows mark syllable types analysed in panel C. **C**. Three example phrase types of bird 3 appear in different probabilities in the 1^st^, 2^nd^, and 3^rd^ thirds of the song in which they are produced. **D**. Histograms showing distributions of syllable repeat numbers in 5 different example phrase types of bird 4. **E**. Pie showing the phrase transition outcome probabilities following the dark red phrase in panel A. Light and dark blue pie slices mark the transition outcomes in panel A. Gray marks other outcomes. **F**. The transition after the dark red phrase in panel A follows a long-range syntax rule. Pies show the transition outcome probabilities following the two different song histories in panel A, marked by colored boxes - demonstrating a strong dependence on the identity of the 2^nd^ phrase preceding the transition. N-values show numbers of history conditioned transitions in each context.

To characterize the changes in canary syntax rules, we analyzed both acoustic and syntactic properties of songs over time periods of 1-3 weeks. We found syntax rules that diverge across days in a range of sequence context dependencies - rules that depend on the context of the syllable identity alone, the context of the preceding phrase, and up to the order and identity of the 2-7 phrases that precede a transition. We found a similar process occurring in the acoustic properties of syllables. Interestingly, within individual birds most changes, acoustic and syntactic, took a similar form of either an increase or a decrease in variability.

These findings suggest that canaries in their mating season undergo processes that manifest in a global move between order and disorder. This process may be part of these birds’ annual cycle of song plasticity [4] and establish canaries as an animal model for studying neural mechanisms that underlie naturally-dynamic syntax rules.

## 2 Results

We recorded songs of six canaries individually housed in soundproof chambers over 1-3 weeks. We leveraged our recently-developed deep learning algorithms [45, 46, 51] that afforded annotating the timing and identity of syllables in more than twenty five thousand canary songs - an order of magnitude more than previously done [31, 45, 39]. The birds’ individual repertoires included 20–47 different syllables with typical durations of 7–330ms. The average number of syllable repeats per phrase type ranged from 1 to 25 (0.05 and 0.95 quantiles across syllable types of 6 birds), with extreme cases of individual phrases exceeding 10 seconds and 130 syllables (Fig. 1D).

Across birds, phrase transitions that follow 75-95% of the phrase types had multiple possible outcomes in different songs. Transitions that follow the other phrase types were deterministic - always leading to the same outcome. Across all birds, transitions following 17-35% of the phrase types depended on long-range song history (Fig. 1E,F). Their outcome had significant statistical dependence on phrase transitions that preceded the currently produced phrase. This is a higher percentage than we previously reported, 15% in [39], owing to the much larger dataset that allowed reliably estimating the probability of rare phrase sequences.

Changes in canary song syntax may occur in multiple hierarchical levels: syllables, phrases, and the long range dependencies in phrase sequences. To characterize syntax dynamics, we quantified these syntax properties in subsets of songs that were recorded in different days.

### 2.1 Canaries change the context and frequency in which they produce various syllable types

We examined *R*^(*day*)^(*s*), the daily rate by which birds use their phrase types (’*s*’, same as syllable types in this context. See Methods) and found significant changes across days. For example, Fig. 2A shows that bird 3 used 11 out of its 20 phrase types in significantly different rates in days that are a week apart (Bootstrap, *p <* 0.025, see Methods) leading to a significantly different distribution *R*(*s*) (Kolmogorov-Smirnov test. D=0.11, *p <* 0.0001).

**Figure 2:**
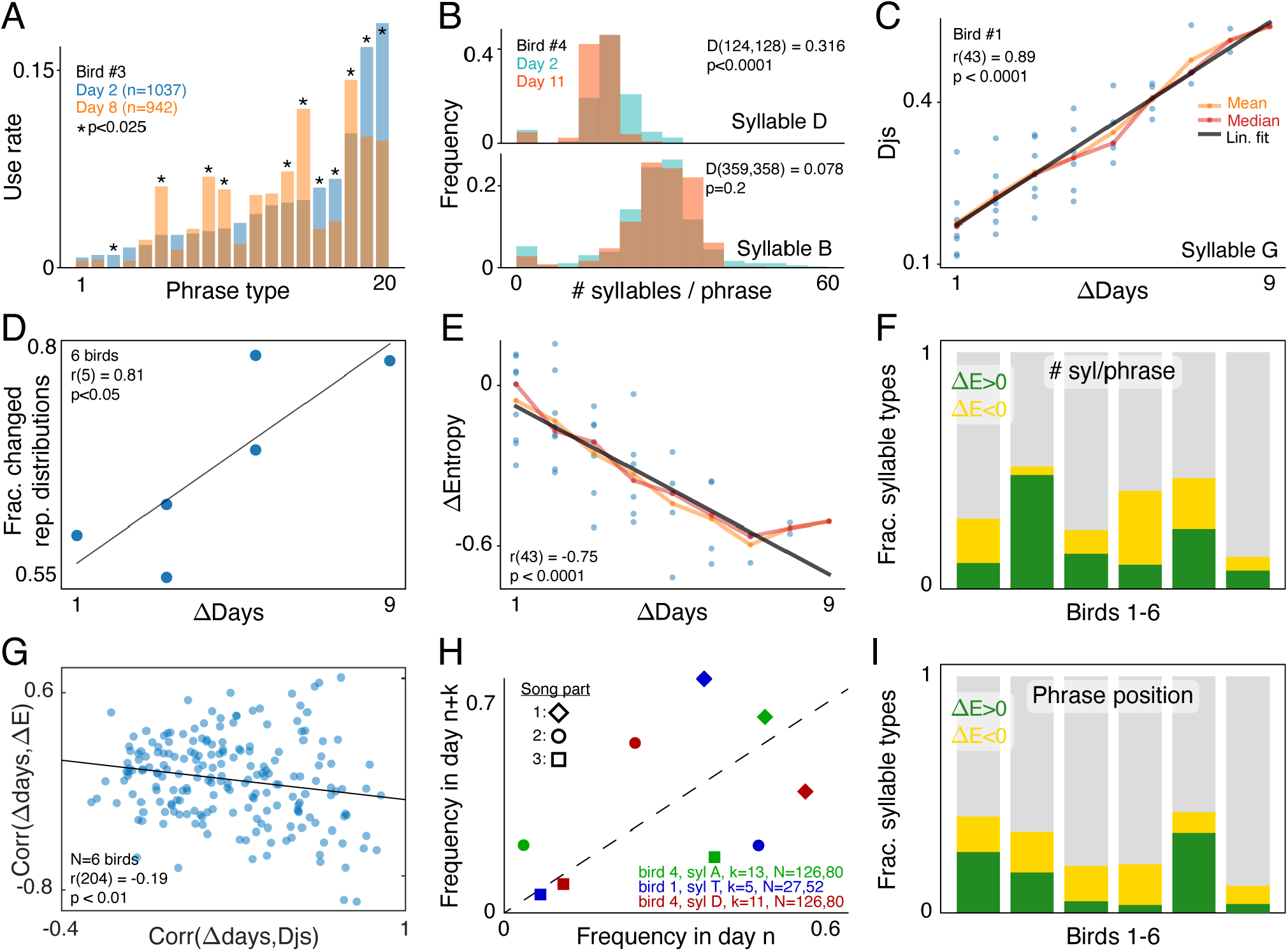
The rate of singing different syllables, their repetition in phrases, and position in song change across days. **A.** Blue bars show bird 3’s rate of using its phrase types in the 2^nd^ recording day (x-axis), *R*^(2)^(*phrase type*). Orange bars show the rates a week later. (★: Bootstrap test, *p <* 0.025). **B**. Repeat distributions, 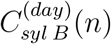 and 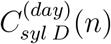 of bird 4’s syllables B,D. Blue and red distributions are calculated in the 2^nd^ and 11^th^ days. KS-tests compare the distributions (D,p values). **C**. Dots show the distance between repeat distributions, 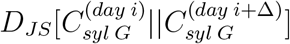 as a functions of Δ (x-axis). Black, orange and red lines show the linear trend (Pearon’s r,p), mean and median. **D**. Markers show fractions of syllables that significantly change their repeat distributions in 6 birds. Fractions increase with the gap (x-axis) between the first 3 days and the last 3 days of recording (Pearson’s r,p). **E**. Dots show the entropy difference between bird 1’s syllable G repeat distributions (same as panel C), 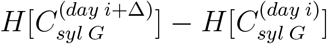 as a functions of Δ (x-axis). Black, orange and red lines show the linear trend (Pearon’s r,p), mean and median. **F**. Green (yellow) bars show the fractions of syllable types with significant positive (negative) Pearson correlation between the repeat distribution entropy difference, 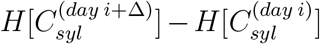 and the gap between days, Δ. **G**. X-axis shows the correlation between the day gap and the distance between repeat distributions. Y-axis shows the correlation between the day gap and the difference between the entropy of repeat distributions. Dots show all syllables of 6 birds. Line shows the trend (Pearson’s r,p). **H**. Examples of the probability of 3 phrase types (colors) to appear in each third of the song sequence (marker shape) is measured in two different days (x,y axes). Dashed line shows the unity. **I**. Same as panel F, but for the distribution of phrase position in song, 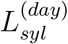

The difference between daily-estimated *R*(*s*) distributions can depend on the gap between days. To test this possibility, we used the Jensen-Shannon divergence to quantify the difference between distributions. For example, in Fig. 2A, the distance between the distributions we calculated in the 2^nd^ and 8^th^ recording days is *D*_*JS*_[*R*^(2)^||*R*^(8)^] = 0.25. We checked if longer gaps between days predict larger differences between *R* distributions by calculating the Pearson correlation between the inter-day gap, Δ*days* = *day*_2_ − *day*_1_, and the distance metric 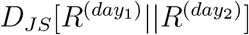 This analysis revealed significant positive correlations in all six birds (Pearson’s *r* = 0.41−0.9, *p <* 0.05, Supp. Fig. 1) indicating that distributions move apart across days.

The movement of these distributions can lead to a more, or less, random use of syllables. The most random case is the flat distribution in which all syllables have the same rate. This distribution has the maximal entropy (see Methods) *H*[*R*^(*d*)^]. The least random case, whose entropy is zero, is when only one syllable type is used and all other syllable types are not used. We tested if the gap between days correlates to the entropy difference. Out of six birds in our study, one had a positive correlation, two had a negative correlation and three did not have a significant correlation (data not shown).

Together, these results suggest that canaries change the rate by which they use syllable types and that only some birds use syllables more or less randomly over days.

A different picture emerged when we examined distributions of syllable repeat numbers, 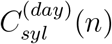 (c.f. examples in Fig.1D). Comparing across days revealed, in each bird, syllables whose repeat distributions changed and others that didn’t (c.f. Fig. 2B). We tested if the gap between days correlated to the distance between repeat distributions and found significant correlations in all birds (25%-89% of all syllable types in birds 1-6 showed significant Pearson correlations, e.g. Fig. 2C, Supp. Fig. 2). To group birds we estimated *C*_*syl*_(*n*) in data from the first three days and the last three days of each bird’s recording. Fig. 2D shows that birds that were recorded for more days also had more repeat distributions that changed.

To test if repeat distributions become more or less random in time we calculated their entropy difference, 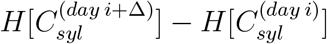 as we did for *R*(*s*) above, and found that it correlates to the day gap, Δ, in 0.1-0.5 of the syllable types (Fig. 2E,F). In some birds most of the repeat distribution lowered their entropy - the repeat numbers became more stereotyped - and in other birds it was the opposite. Across 6 birds, repeat distributions that moved apart across days tended to also decrease in entropy (Fig. 2G, Pearson’s r=-0.19, *p <* 0.01, shows the relation between two correlations: The correlations of the day gap and the repeat distributions’ distance (*D*_*JS*_), and the correlations of the day gaps to the change in repeat distribution entropy).

Apart from the rate of using phrases and the syllable repeats in phrases, we also checked the probabilities for phrases to appear in different parts of song. To do that, we divided each song into three equal parts and calculated, 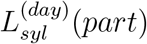 the probability for phrase type ’syl’ to appear in parts 1/3, 2/3, 3/3 (for example in Fig. 1C). Fig. 2H shows 3 example phrases that changed these probabilities. Across all phrase types, as we did with the distributions of syllable repeats, we examined the relations between gaps of days and the distance between the daily-measured distributions 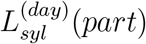 We found significant correlations in all birds (15%-59% of all phrase types in birds 1-6, Supp. Fig. 2). To test if these distributions also changed their randomness we tested for entropy changes, 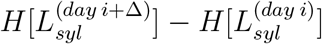 and their correlations to the day difference Δ. Fig. 2I shows that, like the repeat distributions, in some birds most of phrase location distribution lowered their entropy - the position in song became more stereotyped - and in other birds it was the opposite.

### 2.2 Canaries change transition probabilities and variability across days

In the previous section we observed that canaries change the probability of using the different syllables in their repertoire depending on *zero order* contexts - rate, position, and repeats of each syllable type irrespective of other syllable types. In this section we turn to the *higher order* syntactic dependencies of canary phrase sequences - to phrase transitions that depend on the order and identity of the phrase types that preceded them [31].

Fig. 3A illustrates a transition following phrase ’a’. We measure the transition’s outcome probabilities, for example the probability to transition from phrase ’a’ to phrase ’b’, *p*(*b*|*a*) ≡ *P*_*a*_(*b*), disregarding phrase types that preceded ’a’. For brevity, we refer to transition outcome probabilities that are measured irrespective of farther upstream elements as 1^st^ order transitions.

**Figure 3:**
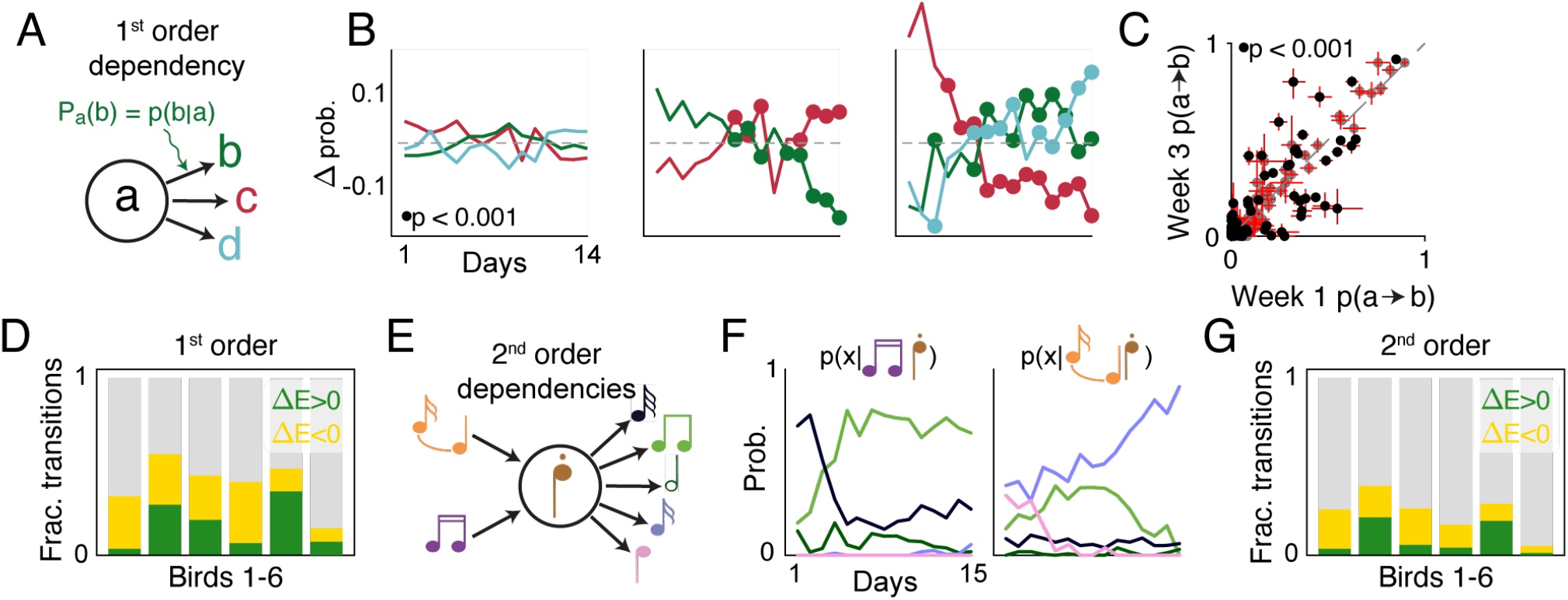
Canary transitions with 1^st^ and 2^nd^ order dependencies on the preceding phrase sequence change across days. **A.** Schematic of a 1^st^ order transition - a branch point in which phrase ‘a’ is followed by one of several other phrases (Here, for example, ‘b’,’c’, or ‘d’). **B**. Transition probabilities following 3 specific phrases (‘a’ in the panel A) to target phrases (colors, different phrase types in each panel) change or remain stable across days. Marker show significant change from the first day (Bootstrap, *p <* 0.001). **C**. Probabilities of all specific pairwise transitions of one bird calculated in the 1^st^ and 3^rd^ week (axes). Error-bars show the 0.05 and 0.95 CL. Full circles show probabilities that changed significantly (Bootstrap, *p <* 0.001). **D**. Green (yellow) bars show the fractions 1^st^ order transitions, defined by the phrase types that precede them, with significant positive (negative) Pearson correlation (*p <* 0.025) between their outcome entropy difference, 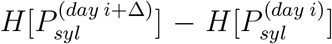 and the gap between days, Δ. **E**. Schematic of an example 2^nd^ order transition in which the phrase marked by the brown note transitions to one of 5 possible outcomes and the probabilities of these outcomes also depend on the identity of the 2^nd^ preceding phrase. **F**. Line colors match the transitions outcomes from E. The two panels show the change in these outcome probabilities across days, 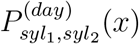 separated by the identity of the 2^nd^ preceding phrase. **G**. Same as D but for the 2^nd^ order transitions, 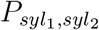 defined by pairs of phrase types that precede them.

In Bengalese finches, 1^st^ order transitions were shown to be actively maintained (keep their value for months and return to baseline when perturbed [29]). Likewise, we found that some canary 1^st^ order transitions did not change across 2-3 weeks, but many others did. Fig.3B,C show individual transition probabilities that changed by 10-80% across 2 weeks. We used the Jensen-Shannon divergence to quantify the change in the 1^st^ order transition distributions 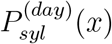 the probabilities for transitioning from phrase ‘syl’ to phrase ‘x’. We also estimated the entropy differences, 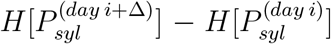 and its relation to the difference between days, Δ. Much like the distribution of syllable repeats, the transition distributions following 35%-65% of the phrase types in birds 1-6 grow farther apart across days (Supp. Fig. 3). Similarly, a significant fraction of daily-estimated transition distributions changed their entropy in correlation to the difference between days (Fig. 3D, Supp. Fig. 4 shows example ΔEntropy vs ΔDays scatter plots).

**Figure 4:**
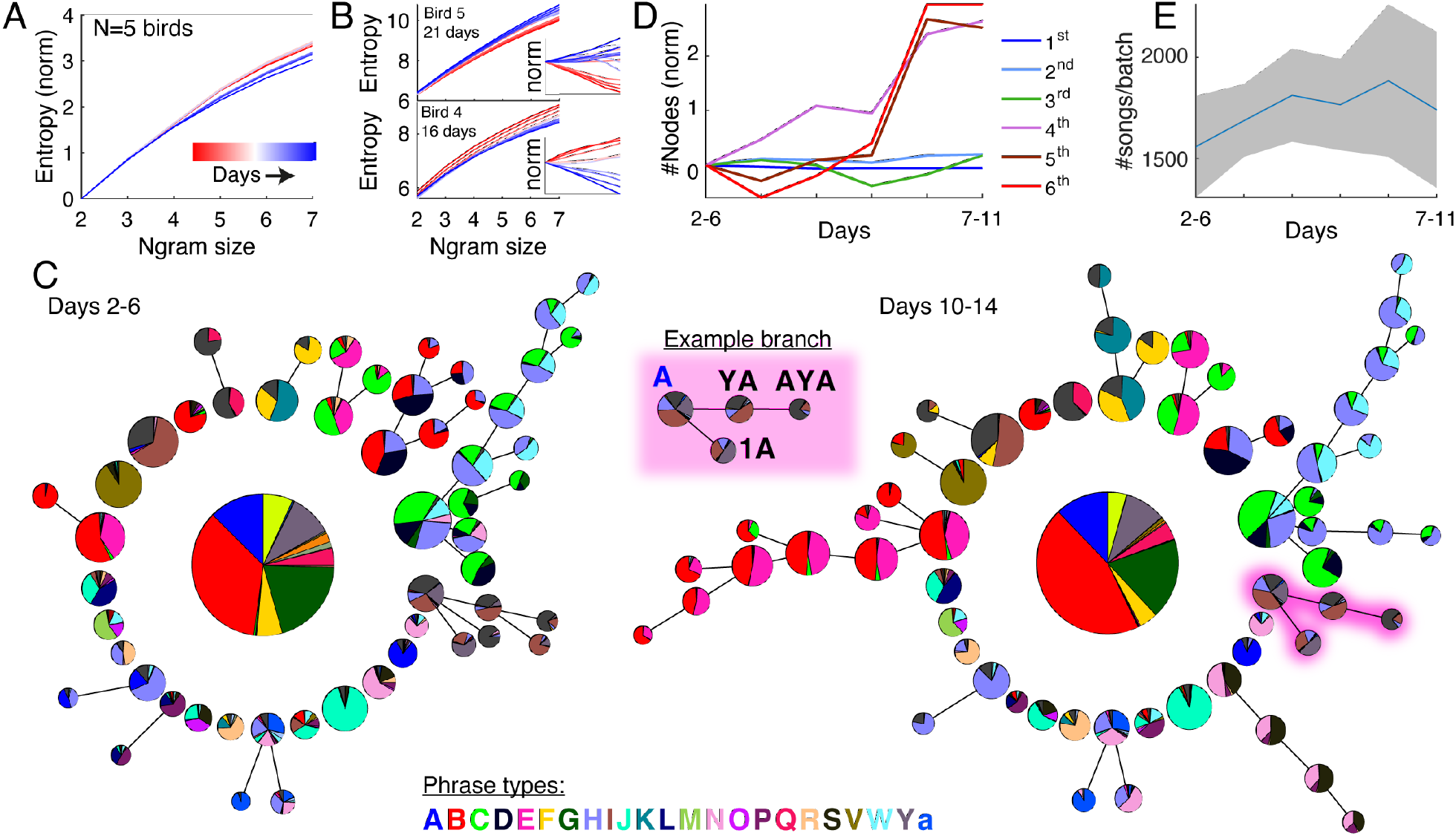
Ngram entropies and Probabilistic Suffix Trees reveal a directional change in the long-range order of canary song. **A.** Ngram entropy is the entropy of sequences of N consecutive phrases (x-axis, Methods). Lines show values after subtracting the entropy for N=2 and averaging across 5 birds. Colors annotate batches of 5 days starting on the 2^nd^-11^th^ day. **B**. Top, the Ngram entropy (y-axis, not normalized) of bird 5, recorded for 21 days, increases for all Ngram sizes (x-axis. Colors, red to blue, mark the days). Bottom, bird 4, recorded for 16 days, shows the opposite trend. Insets show the values normalized by subtracting the mean across days. **C**. Probability Suffix Trees (PSTs, Methods) are graphical models that capture the dependence of canary phrase transitions on song history. Each branch captures the transition from a specific phrase type. Pies describe transition probabilities following specific song histories, annotated for example in the highlighted ’A’ branch in the right PST (supfig shows the fully annotated graph). This branch is of depth 3 and indicates a that the transition following phrase ’A’ is statistically dependent on transitions 3 steps prior. **D**. The number of PST nodes is normalized for each bird to account for the size of the birds’ individual repertoires (see Methods). Lines show the fractional change compared to the first batch of days (days 2-6) averaged across 5 birds. Colors indicate the depth of nodes. **E**. Line and shaded area show the mean and SE of the number of songs per batch of days across the 5 birds that were used for creating panels A-D.

A transition’s entropy reflects its variability. In a low entropy transition, one outcome is far more likely than others (in an extreme case the entropy is 0 and the transition is deterministic). In contrast, outcomes of a high entropy transition have similar probabilities. In some of the birds we found more 1^st^ order transitions that became less variable across days and fewer 1^st^ order transitions that became more variable (e.g. birds 1,3,4 in Fig. 3D). In other birds, for example bird 5 in Fig. 3D, we found the opposite trend.

A canary’s phrase transition is *strictly 1*^*st*^ *order*, or Markov, if it depends only on the currently produced phrase. In six birds, 65-83% of phrase types preceded Markov transitions. The transitions that follow the rest of the phrases depended on deeper song history. These phrase transitions can, in principle, follow different - even opposite - changes across days than the 1^st^ order transitions. To test this possibility, we first examined 2^nd^ order transitions – the daily estimated probabilities 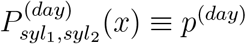 of transitioning to phrase *x* after singing the pair of phrases *syl*_1_ → *syl*_2_. For example, Fig. 3E shows a phrase type, marked as a brown musical note, that was produced in different songs following two possible upstream phrases and preceding five possible downstream phrase types (marked by colored notes). Fig. 3F shows the 2^nd^ order transition probabilities. These probabilities change across days and also depend on the identity of *syl*_1_ (purple or orange notes in this example, *syl*_2_ is the brown note).

As we did for 1^st^ order transitions, we used the Jensen-Shannon divergence to quantify the changes in 2^nd^ order transition distributions, 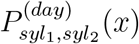 and estimated their entropy differences, 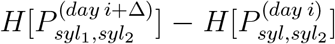 across Δ days. We found that transition distributions following 12%-35% of the phrase types in birds 1-6 grow farther apart across days (Supp. Fig. 3). Similar to the 1^st^ order analysis above, a significant fraction of daily-estimated 2^nd^ order transitions changed their entropy in correlation to the difference between days (Fig. 3G, Supp. Fig. 5 shows example ΔEntropy vs ΔDays scatter plots). In all birds, the fraction of 2^nd^ order transitions with days-correlated entropy changes were smaller than the fractions of 1^st^ order transitions with days-correlated entropy changes. But the same birds that had mostly negative correlations for 1^st^ order transitions (Fig. 3D), also had mostly negative correlations for 2^nd^ order transitions (Fig. 3G).

**Figure 5:**
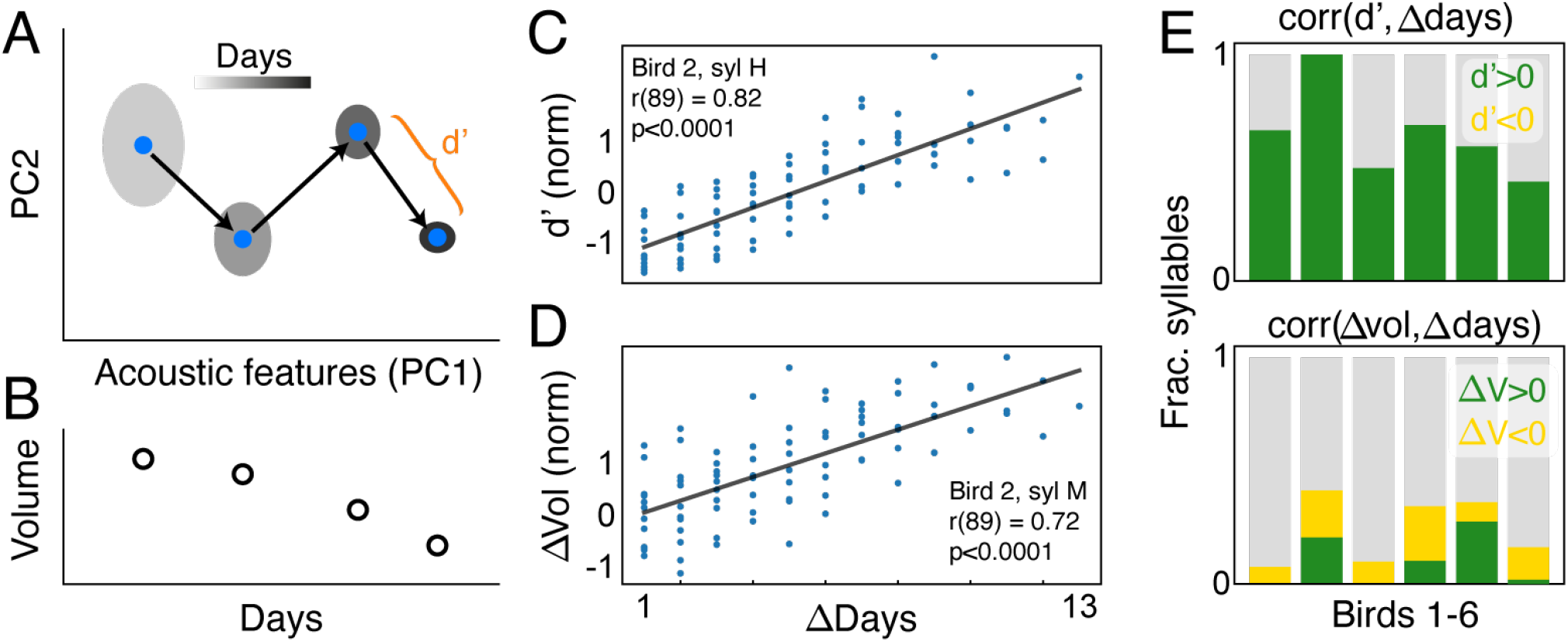
Canary syllable acoustics change across days. **A.** An illustration showing daily measures of a specific syllable’s acoustics. Syllable renditions form a cloud of points in an 7-dimensional acoustic features space (ellipses). From each cloud we extract the mean, *µ*_*syl*_(*day*) (blue dots) and covariance matrix, Σ_*syl*_(*day*). These measures are used to calculate *d*^*′*^(*day*1, *day*2), the difference between days using the Mahalanobis distance. **B**. For each day we measure the volume taken by the syllable renditions in A (see Methods). **C**. Dots show the distance 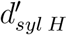 (y-axis) strongly correlated to the day difference Δ (x-axis) for an example syllable (syllable H of bird 2, Pearson’s r,p). **D**. Dots show the volume difference Vol_*syl M*_ (*day i*) − Vol_*syl M*_ (*day i* + Δ) (y-axis) strongly correlated to the day difference Δ (x-axis) for an example syllable (syllable M of bird 2, Pearson’s r,p). **E**. Green (yellow) bars show fractions of syllable types for which significant positive (negative) correlations (Pearson, *p* < 0.025) were found between the day difference (x-axis in panels C,D) and the acoustics difference (top) or volume in acoustics space (bottom).

The similarity between the fractions of 1^st^ and 2^nd^ order transitions with same-sign entropy changes across days is not surprising since many canary transitions are Markov. Still, we identified syllable types that precede 1^st^ order transitions and 2^nd^ order transitions whose entropy change across days was unrelated, and even opposite. Specifically, in every bird we found syllables *syl*_1_, *syl*_2_ for which the change in entropy of 2^nd^ order distribution, 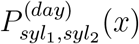 correlated to the difference in days, and the change in entropy of the 1^st^ order distribution, 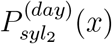, either had no correlation to the difference in days or even had the opposite correlation. Across all 6 birds, these opposite trends occurred in 23/204 syllable types (≈ 11.25%, Pearson’s *p* < 0.025)

Beyond 2^nd^ order syntactic dependencies, some canary phrase transitions were shown to depend on 5-7 transitions upstream ([31, 45]). These song history dependencies may also change across days and weeks and implicate a global process that affects a canary’s song syntax rules.

### 2.3 Canary syntax moves between order and disorder

The above sections demonstrated that for several canaries more 1^*st*^ and 2^*nd*^ order transitions became less variable across days compared to transitions that became more variables. Other canaries demonstrated the opposite trend. More of their 1^*st*^ and 2^*nd*^ order transitions became more variable.

To better understand the process of syntax rule changes we divided each canary’s songs to batches of 5 consecutive days with 4 days overlap. For example, if one batch contained songs from recording days 2-6, the next batch was from days 3-7. This division resulted in an average of between 1500-2000 songs per batch per bird (Fig. 4E). In each batch we examined the distribution of phrase sequences of length *N* = 2 − 7. If a canary produces every *N* -long phrase sequence, or Ngram, with equal probability, it means that the bird’s syntax is maximally disorganized in that *N* ^th^ order.

Canaries do not produce every possible Ngram. To estimate the degree of organization in Ngram orders 2-7 we measured the entropy of the Ngram distributions [31]. Five of our birds were recorded for 11 days or more so we could measure their Ngram entropies in 6-16 consecutive 5-day batches.

Fig. 4A shows the Ngram entropy values averaged across 5 birds. The average values decrease across days, most prominently in the large Ngram sizes. However, averaging masks the trends of individual birds. Specifically, out of 5 birds 2 exhibited a monotonous decrease in Ngram entropy in all orders (Fig. 4B, bottom). In contrast, 2 birds exhibited a monotonous increase across days (Fig. 4B, top).

Observing that Ngram distributions of multiple orders monotonously increase their entropy indicate a global process by which the songs become disorganized. Observing Ngram distributions that decrease their entropy indicates the opposite process - songs that become more regular. To chart this process we estimated the variable-length Markov chains that capture the song history dependencies of phrase transitions [31, 45].

This syntax structure is captured parsimoniously by probabilistic suffix trees (PSTs[10]). The root node in these graphical models, appearing in the middle of the graphs in Fig. 4C, represents the zero-order Markov, or base rate, frequencies of the different phrases, labeled in different colors and letters. The branches, organized in a circle and emanating outwards, represent the set of Markov chains that end in the specific phrase types. For example, the pies in the ’A’ branch, highlighted in Fig. 4C, show transition probabilities following phrase ’A’. The pie titles describe dependencies on song history of different depths - a 1-deep dependency ’A’, 2-deep dependencies ’YA’ and ’1A’, and the 3-deep dependency ’AYA’ - and every time the ’A’ phrase occurs, the deepest ’leaf’ that matches A’s upstream history determines the transition outcome probabilities. Together, these dependencies parsimoniously describe the 3^rd^ order Markov chain that captures the transition from phrase ’A’. For presentation simplicity, we removed the node labels from the rest of the branches in Fig. 4C.

For each bird, we estimate the PST in each 5-days batch. We then count the number of nodes of different depths in the tree. To group birds together, we normalize the number of nodes at various depths of the tree by the potential number of nodes (See Methods). The fractional changes in average number of PST nodes at different depths is shown in Fig. 4D. This analysis reveals that the overall entropy decrease, demonstrated in high order Ngrams in Fig. 4A, is also seen as the increase in the number of PST deep branches (nodes of the 4^th^-6^th^ order in Fig. 4C). This dynamics indicates an increase in long-range syntax dependencies. As an example, Fig. 4C shows two PSTs estimated for bird 4. One PST (Fig. 4C, left) is calculated from songs in recording days 2-6 and the other PST (Fig. 4C, right) is calculated for days 10-14. Both the Ngram entropy changes (Fig. 4B) and the dynamics of 1^st^ and 2^nd^ transition entropies suggest that the song of this bird becomes more organized across days. The PSTs in Fig. 4C show that the song organization manifests as more PST nodes of order *>*3 (4 such nodes in days 2-6, all in one branch, and 11 nodes in days 10-14, on 3 branches). Put differently, more of this bird’s transitions, captured by more PST branches, develop long-range dependencies on song history and make the song more organized.

### 2.4 Canaries change the acoustic properties of syllables across days

The previous sections showed that canaries change the probability of singing their different syllables depending on a variety of syntactic contexts. Work in Bengalese finches discovered that syntactic context also impact the acoustic features that define individual syllables [25]. We therefore examined whether canary syllable acoustics exhibits similar changes across days as did their song syntactic properties.

We captured the acoustic features of each syllable rendition using 7 measures defined in [25] (e.g. duration, fundamental frequency, spectral entropy, see Methods). For each measure, we calculated the median value from syllable repeat phrases and normalized across all phrases of the same syllable. Fig. 5A illustrates the resulting daily cloud of 7-dimensional points (projected on two principal components for presentation only) from which we calculated the daily variability - the volume of the cloud Vol_*syl*_(*day*) (Fig. 5B) - and the acoustic difference between days, 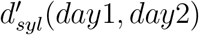 (see Methods).

To test if syllable acoustics changed across days we measured the correlation between the difference in days and the difference in syllable acoustics *d*^*′*^ (Fig. 5C). In 6 birds, 45-100% of the syllables exhibited this correlation (Pearson, *p <* 0.025 Fig. 5E, top). Similarly, we calculated the correlation between the difference in days and the change in syllable acoustic variability (Fig. 5D). This calculation revealed that each of the 6 birds had syllables that changed their acoustic variability. The bottom panel in Fig. 5E shows that in some birds, most significant trends showed a reduction in variability. In others we found the opposite trend.

### 2.5. Canary syntactic and acoustic changes occur in similar trends across days

Thus far, we observed that canaries change various syntactic properties that define the probability of producing syllables. In section 2.3 we focused on 5 canaries that were recorded for 11 days or more and found that most of them exhibited a monotonous move between order and disorder. In the above section we found a similar trend in the syllables’ acoustic variability, measured as the syllable renditions’ volume in acoustics space (Fig. 5E, bottom).

To begin charting the relations between the syntactic and acoustic properties, we compared the fractions of various changes that occur in correlation with the differences of days. Fig. 6A shows these fractions for each bird and separates positive and negative correlations (*ρ >* 0 and *ρ <* 0). This analysis summarizes the results of previous sections (Fig. 2F,I, Fig. 3D,G, and Fig. 5E) and reveals that, in four out of six birds, most measures show the same trend. In bird 4, all five measures, syntactic and acoustics, show more variability decrease across days, and in bird 5 we find the opposite - all measures show variability increase. In two more canaries, birds 1 and 3, we find that all but one measure have the same trend and the acoustic variability is in agreement with the majority of the syntactic variability. In the last two birds, including bird 6 that was only recorded for a week, there is no clear global trend but the acoustic variability has a similar trend to most syntactic variabilities.

**Figure 6:**
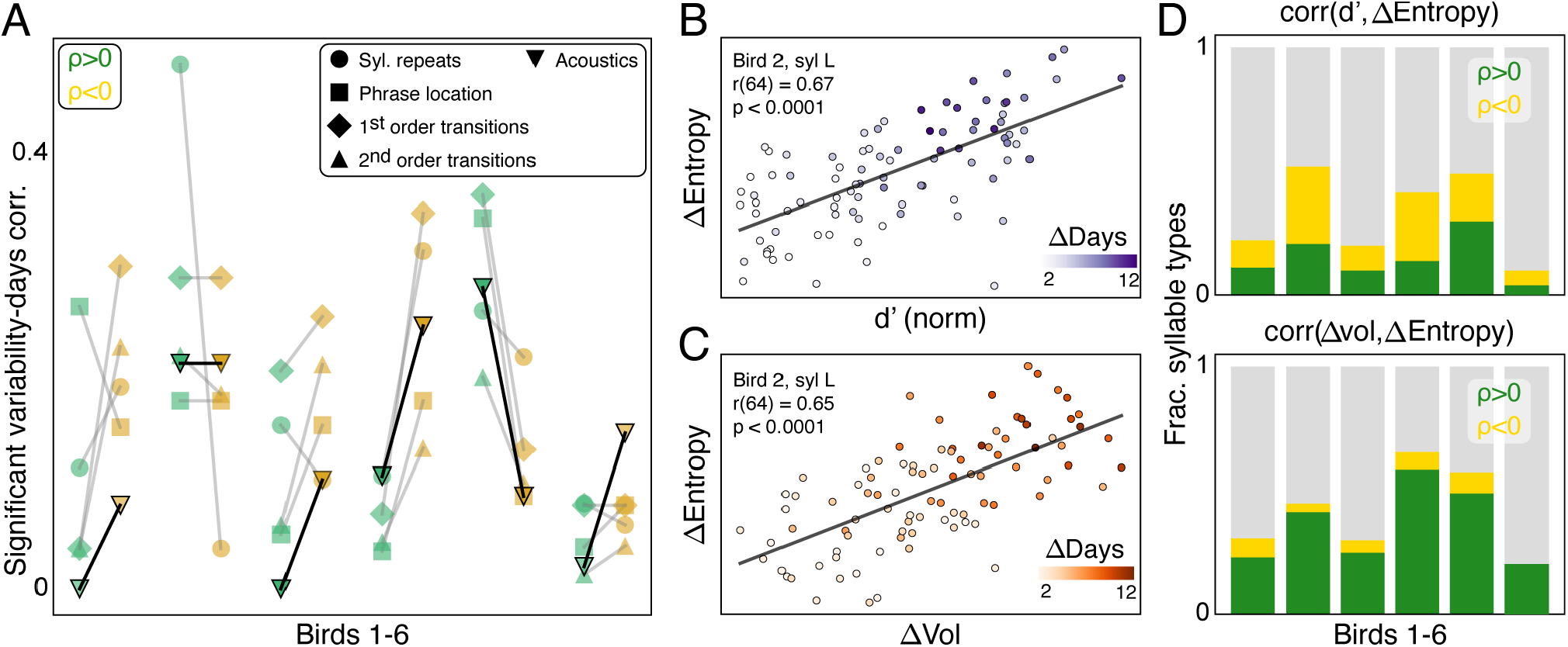
The variability of syntactic and acoustic properties exhibit similar changes across days. **A.** Summary of results from Fig. 2F,I, Fig. 3D,G, and Fig. 5E. Y-axis shows fractions of significant correlation tests between the difference in days and the change in the variability of five syntactic and acoustic properties (Marker shapes). Results are shown for 6 birds (x-axis) and marker colors (green,yellow) separate positive and negative correlations coefficients (*ρ*). Lines connect fractions of positive and negative correlations of the same properties. **B**. Dots mark pairs of days for which we measured a syllable’s acoustic difference (*d*^*′*^, x-axis, syl L of bird 2 in this example) and change in the entropy of the next 1^st^ order transition, *P*_*syl L*_(*x*). Color-bar shows the day-difference, Δ*Days*. **C**. Same as B but instead of the acoustic difference *d*^*′*^, the x-axis shows the difference in acoustic variability, ΔVol. **D**. Green (yellow) bars show fractions of syllable types for which significant positive (negative) correlations (Pearson, *p <* 0.025) were found between the change in 1^st^ order transition entropy and acoustic difference (top panel) or change in acoustic variability (bottom).

These similarities between the dynamics of both acoustic and syntactic properties of canary song can take many forms. We focused on relating syllable acoustics to the variability of the phrase transition that immediately follow them - the 1^st^ order transition. Fig. 6B,C show an example syllable (syl. L of bird 2) for which both the acoustic difference, *d*^*′*^ and variability difference, ΔVol, correlated to the entropy change of the next phrase transition.

Syllable acoustics was previously shown to impact the upcoming transition [15]. The top panel in Fig. 6D shows that, in all birds, as syllable acoustics moved farther apart, the variability of the next transition could increase or decrease. This observation is not surprising and stems from the above findings that acoustic differences only increase across days (Fig. 5E, top). The bottom panel in Fig. 6D shows the fractions of significant correlations between acoustic variability changes and the phrase transition entropy changes. These fractions are much larger, in all birds, than the fractions of significant correlations between the acoustic variability changes and the differences in days (Fig. 5E, bottom). Moreover, in all birds the correlations in Fig. 6D (bottom) are overwhelmingly positive - suggesting a relation between the dynamics of syllable acoustic variability and variability of the transitions that follow these syllables. We examined the possible lag between the dynamics of acoustic variability and the dynamics of transition entropy and found no significant shift at the resolution of days (Wilcoxon signed rank tests the hypotheses that the distribution of lags is shifted from 0, *p* > 0.2, Supp. Fig. 6).

In sum, the above results show that canaries change syntactic and acoustic properties that define their repertoire in the spring mating seasons. In some birds, these changes indicated a global move from order - low variability of syntactic and acoustic properties - to disorder. In other birds we found the complete opposite - a move from disorder to order.

## 3 Discussion

Songbirds are an excellent model for studying the neural mechanisms that underlie vocal learning and skilled production of motor sequences. In landmark studies, neurophysiological and behavioral experiments leveraged the stereotyped song of zebra finches [22, 23, 24, 30, 49, 47, 35, 20, 14] and the crystallized transitions of Bengalese finches [27, 11, 28, 44, 25, 15, 17]. These experiments followed a standard approach in songbird neuroscience research - to record song and neural activity from birds that are housed in soundproof boxes in laboratory settings. Following the same protocols, our results suggest that the spring mating song of canaries is not crystallized.

We examined canary syntax rules that govern the probabilities of producing different syllables depending on various contexts of song sequence - ranging from the syllables’ position in song, the numbers of syllable repeats in trill phrases, to the dependence of phrase transition probabilities on the preceding sequence of 1-7 phrases. We also analyzed the change in canary syllable acoustics across days and the change in acoustic variability. Our overarching observation, summarized in Fig. 6A, is that individual canaries exhibit a directional change in variability across all measures - acoustic and syntactic - suggesting a global move between order and disorder that can be observed across days and weeks of the canary spring mating season.

Unlike stable behavior, which is either crystallized or actively maintained, we cannot rule out the possibility that the behavioral dynamics we observed is influenced by the experimental conditions and time line. In our analyses, we removed the first day song recording and replicated the findings in six canaries recorded across 1-3 weeks in April-June. Our robust observations, repeated across animals and various measures of syntactic and acoustic variability, will allow studying the underlying neural mechanisms. Still, like many laboratory neuroscience experiments, further exploring the effect of social and environmental variables is needed.

For example, past studies revealed that canaries change their song in the presence of male or female conspecifics [38, 9, 12]. Social contexts may reduce the variability in canary song, like the variability reduction in female-directed zebra finch song compared to singing alone [18].

Several of the birds we recorded exhibited a global variability reduction that allowed robustly observing longer-range dependencies of phrase transitions on the song sequence (Fig. 4B-D). Observing a similar effect in social conditions may indicate that long-range syntax rules play a social role. Alternatively, social conditions may drive canaries to increase song variability, as some birds in our experiment did in isolation. In either case, reduction or increase in variability caused by social experiments will indicate which properties of the male song are potentially perceived and evaluated by the females. In establishing a baseline for canary song dynamics, our results make possible future studies of changes in syntax rules that reflect the flexibility required for adapting vocal sequences to different social and environmental contexts.

Changes in syntax rules and syllable acoustics also afford a new window onto studying neural mechanisms for sensory-motor sequence generation. Birdsong is generated by neural circuits homologous to thalamo-cortical and basal ganglia loops [19] and the choice of the next syllable in the sequence was also shown to depend on auditory input and on adaptation to syllable repeats [15, 28, 16]. Several neural network models were suggested as mechanisms for producing birdsong syllables and transitioning between them (e.g. [40, 21]). Studying these models’ ability to explain syntax rule dynamics will further constraint the possible mechanisms and has the potential to expose neural plasticity associated with learning and modifying motor sequences. Specifically, plasticity of ongoing auditory-motor processing will need to account for the observed global process of moving between order and disorder. Such a process may be part of the canary seasonal cycle of song plasticity and neurogenesis and rely on both neuromodulatory and hormonal changes [37, 36]. Our results establish canaries as a suitable animal model for studying such mechanisms for syntax rule dynamics since most of our test subjects strongly showed the process of moving between order and disorder. Still, individuals showed divergent trends. This individual variability observed among canaries, with some moving towards more stereotyped song and others becoming more variable, may depend on a-priori differences in repertoire, on age, or on other, currently unknown, individual traits. Building on our results, further research may relate individual differences to potential behavioral traits and mating strategies.

Finally, the probabilistic framework of syntactic dependencies is the key advance in statistical language models - a rapidly expanding field of basic and applied research [52]. After being trained on enormous amounts of textual data, these models mimic the context-sensitive probability of word choice in language. Lessons learned from biological systems that learn, produce, and augment syntax rules in rich behavioral repertoires, could both improve the expansive process of training artificial language models or even inform the development of improved language models, particularly those requiring adaptive and context-sensitive capabilities.

## 4 Methods

### 4.1 Data collection and preparation

In this work we analyze data that was previously published in [45]. Songs were recorded from 6 American Singer canaries at Boston University in 2018 and 2019. In the current project we add to the data set by annotating more songs of one of the birds. This data is made public in url.url.

We prepare the data using the same pipeline as in our previous work [45]. Here, we briefly describe the data collection and preparation procedures.

#### 4.1.1. Song screening

Birds were individually housed in soundproof boxes and recorded for 3–5 days (Audio-Technica AT831B Lavalier Condenser Microphone, M-Audio Octane amplifiers, HDSPe RayDAT sound card and VOS Games’ Boom Recorder software on a Mac Pro desktop computer). In-house software was used to detect and save only sound segments that contained vocalizations. These recordings were used to select subjects that were copious singers (≥ 50 songs per day) and produced at least 10 different types of syllables.

#### 4.1.2. Song recording

Data were collected from n=6 adult male canaries between late April and early June of 2018 and 2019 (Audio-Technica AT831B Lavalier Condenser Microphone, M-Audio M-Track-8 amplifiers, and VOS Games’ Boom Recorder software on a Mac Pro desktop computer). Birds were individually housed for the entire duration of the experiment and kept on a light–dark cycle matching the daylight cycle in Boston (42.3601° N) with unlimited access to food and water. None of the birds were recorded in both years. The sample sizes in this study are similar to sample sizes used in the field. The birds were not used in any other experiments. This study did not include experimental groups and did not require blinding or randomization.

#### 4.1.3 Audio processing

##### Song annotation

Song syllables were segmented and annotated by a semi-automatic process. First, a set of 100 songs was manually annotated using a GUI developed in-house (https://github.com/yardencsGitHub/BirdSongBout/tree/master/helpers/GUI). This set was chosen to include all potential syllable types as well as cage noises. The manually labelled set was then used to train a deep learning algorithm (‘TweetyNet’ [45]). The trained algorithm annotated the rest of the data and its results were manually verified and corrected by two expert human annotators in sequence (first in [45] and a second time, verifying and correcting extremely rare errors in preparing the current manuscript). In both the training phase of TweetyNet and the prediction phase for new annotations, data were fed to TweetyNet in segments of 1 s and the output of TweetyNet was the most likely label for each 2.7-ms time bin in the recording.

##### Assuring the separation of syllable classes

The manual steps in the pipeline described above can still miss rare syllable types or mislabel syllables into the wrong classes because of the human annotator’s mistake or because some annotators are more likely to lump or split syllable classes. To address this potential variability in canaries, where each bird can have as many as 50 different syllables, we made sure two annotators agree on the definition of the syllable classes. Then, to make sure that the syllable classes are well separated, all the spectrograms of every instance of every syllable, as segmented in the previous section, were zero-padded to the same duration. An outlier detection algorithm (IsolationForest: https://scikit-learn.org/stable/modules/generated/sklearn.ensemble.IsolationForest.html) was used to flag and recheck potential mislabeled syllables or previously unidentified syllable classes.

### 4.1 Data analysis

#### 4.2.1 Syllable acoustics

##### Acoustic features

Following commonly used definitions of birdsong syllable acoustics (e.g. in [25, 17]), we used a set of 7 features to characterize a syllable. Briefly, for every syllable in our data set, we measured the duration, fundamental frequency, time to half-peak amplitude, frequency slope, amplitude slope, spectral entropy, and temporal entropy (see [25]) - yielding 7 quantities.

These quantities vary in different ranges. For example, the fundamental frequency ranges between 100s and 1000s of Hz while the spectral entropy is of orders 1-10. We therefore normalized (z-scored) the values for each feature separately such that the mean and standard deviation across all syllables is 0 and 1 respectively.

##### Syllable type metrics

We represented each syllable as a point in the 7-dimensional space defined by acoustic features. For each syllable type, *s*, we calculated the mean, *µ*_*s*_(*day*) across all points in a given day (the centroid defining the typical syllable of that day) and Σ_*s*_(*day*) is the daily-estimated 7 × 7 covariance matrix.

##### Acoustic differences between days

To compare renditions of a syllable type between two recording days we used the discriminability, calculated as the Mahalanobis distance:

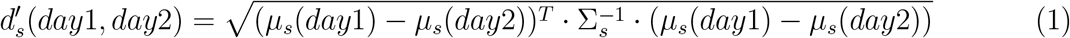

where Σ_*s*_ = (Σ_*s*_(*day*1) + Σ_*s*_(*day*2))*/*2.

To estimate the significance of *d*^*′*^ values we perform bootstrap shuffles, mixing the ‘day’ label of syllables but keeping the number of syllables for each day fixed. Repeated shuffles are then used to create a surrogate distribution of 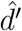 values which we compare to the actual *d*^*′*^ and extract a p-value - the fraction of 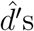 s that are larger than *d*^*′*^.

##### Volume in acoustic space

The volume of an *n*-dimensional ellipsoid, which represents the spread of data in the acoustic feature space, was computed using the covariance matrix Σ of the data points. Prior to this, the data for each syllable was normalized separately by calculating the z-scores for each measurement. This was done by subtracting the mean and dividing by the standard deviation, both of which were computed across all days. The volume *V* of the ellipsoid is given by the formula:

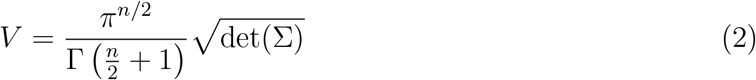

where Γ(·) represents the Gamma function, and *n* is the number of dimensions. The determinant of the covariance matrix, det(Σ), reflects the scaling of the volume based on the spread of the data in each dimension.

#### 4.2.2 Zero order statistics

We use the term “Zero order” to describe statistics of syllable type properties calculated irrespective of other syntax properties defining the context in which syllables appear.

##### Probability to use a syllable in song

In Fig. 1B we show the probability that syllable types are used per song. This probability is different than the frequency by which different phrase types appear in a dataset (described in the next paragraph). For example, if a phrase type appears twice in the same song it will not change the probability per song and it will change the frequency per phrase.

##### Phrase type use rate

To measure the use-rate of a phrase type, *R*(*s*), we first count the number of phrases in which each syllable type was used and then normalize these across all syllable types (so ∑_*s*_ *R*(*s*) = 1). The result of this normalization is a probability distribution of syllable-use rate that is sometimes referred to as the 0th order Markov property of syllable types. We use the Jensen-Shannon divergence to calculate the difference between *R*(*s*) estimated in different days *d*1 and *d*2 (or groups of days). We calculate *H*[*R*], the entropy of *R*(*s*), as a measure of randomness and define,

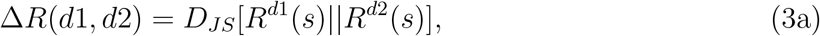

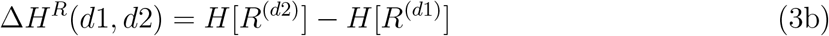

##### Syllable repeats in phrases

For each syllable type, *s*, we calculate the histogram of repeat counts across phrases. We normalize the histogram and create a probability distribution *C*_*s*_(*n*), where *n* is the number of times a syllable is being repeated and ∑_*n*_ *C*^*s*^(*n*) = 1, ∀*s*. Similar to *R*(*s*) above, we define 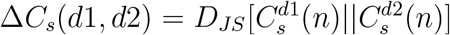 for each *s*. We also calculate *H*[*C*_*s*_] and define 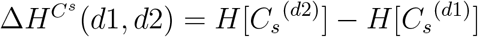

##### Median syllable duration in phrases

For each syllable type, *s*, we calculate the histogram of median syllable duration across phrases. We build the histogram using 25 time bins, *t* and normalize the it to create a probability distribution *T*^*s*^(*t*) (so, ∑_*t*_*T*^*s*^(*t*) = 1, ∀*s*). We calculate Δ*T*^*s*^ (*d*1, *d*2), 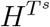, and 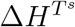 as for *C*^*s*^ above.

##### Phrase position in song

We divided each song into a specified number of parts (in this case, three: start, middle, and end), and calculated 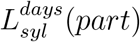 the probability of each phrase type ‘syl’ to appear in these parts. The division of songs into parts was performed using a method that evenly distributes the phrases across the defined sections. Specifically, the number of phrases in the song was divided as evenly as possible among the three parts, ensuring that each part contained a similar number of phrases, with any remainder phrases distributed starting from the first part. For each time period, the relative frequencies of phrases within these divided sections were determined. These frequencies were then normalized to produce probability distributions, reflecting the likelihood of each phrase occurring in different parts of the song.

#### 4.2.3 First order dependencies

We use the term “First order” to describe statistics of transitions between syllable types calculated irrespective of other syntax properties defining the context in which the transition occurs. Transitions between distinct canary syllable types (e.g. *a* and *b*) define transitions between the corresponding phrases (e.g. *A* and *B*). We therefore only refer to first, and higher order, dependencies as phrase transition events.

##### Pairwise transitions

For each transition between distinct and specific phrase types, *A* and *B* s.t. *A* ≠*B*, we calculate *P*_*A*_(*B*) ≡ *P*_*A*→*B*_ = *p*(*B*|*A*), the probability for *B* to follow *A* in a song. We consider a transition to be deterministic if its probability is *P*_*A*→*B*_ ≥ 0.95. We estimate the confidence levels for each *P*_*A*→*B*_ by first calculating *P*_*A*_(*x*) = *p*(*x*|*A*) where *x* are all the phrase types and the end of song event. Then, using *N*_*A*_, the measured number of times phrase *A* occurred, we create *N*_*bootstrap*_ = 10, 000 samples of *N*_*A*_ phrase types drawn i.i.d. from *P*_*A*_ (*x*). We convert each sample to a probability distribution creating a set 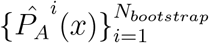 From this set we calculate the 0.025 and 0.975 quantiles for the phrase type *B*.

When comparing song corpora recorded in different days *d*1 and *d*2 (or groups of days) we calculate the difference 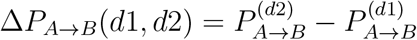

##### First order transition points

For each phrase type *A* we calculate the distribution function *P*_*A*_(*x*) = *p*(*x*|*A*). We calculate the entropy *H*[*P*_*A*_] as a measure of randomness. Given two days (or groups of days) *d*1 and *d*2, we calculate 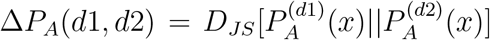 and 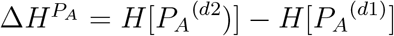

#### 4.2.4 Second order dependencies

We use the term “Second order” to describe statistics of transitions between syllable types calculated in the context defined by the identity of a third syllable type.

##### Triple-wise transitions

For each triplet of phrase distinct types, *A, B*, and *C*, we calculate *P*_*AB*_(*C*) ≡ *P*_*B*→*C*|*A*_ = *p*(*C*|*AB*) ≡ *p*(*A* → *B* → *C*|*A* → *B*), the probability for *C* to follow *AB* in the sequence of song phrases. We estimate the confidence levels for each *P*_*B*→*C*|*A*_ by first calculating *P*_*AB*_(*x*) = *p*(*x*|*AB*) where *x* are all the phrase types and the end of song event. Then, using *N*_*AB*_, the measured number of times phrase sequence *AB* occurred, we create *N*_*bootstrap*_ = 10, 000 samples of *N*_*AB*_ phrase types drawn i.i.d. from *P*_*AB*_(*x*). We convert each sample to a probability distribution creating a set 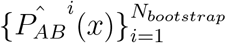 From this set we calculate the 0.025 and 0.975 quantiles for the phrase type *C*.

##### Second order transition points

For each phrase sequence *AB* we calculate the distribution function *P*_*AB*_(*x*) = *p*(*x*|*AB*). We calculate the entropy *H*[*P*_*AB*_] as a measure of randomness. Given two days (or groups of days) *d*1 and *d*2, we calculate 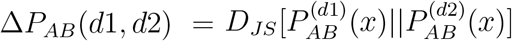 and 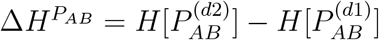

##### Analysis of difference between group of days

Given two groups of days, *d*1 and *d*2, we calculate quantities Δ*R*, Δ*H*^*R*^, Δ*C*, Δ*H*^*C*^, Δ*T*, Δ*H*^*T*^, Δ*P*, Δ*H*^*P*^ (here omitting the indices *s, A, AB, d*1, and *d*2 for brevity). To estimate the significance of each between-group difference values we perform bootstrap shuffles, mixing the group label (*d*1, *d*2) of songs but keeping the number of songs for each group fixed. For example, repeated shuffles create a surrogate distribution of 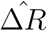 values which we compare to the actual value Δ*R* and extract a p-value - the fraction of 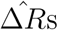 that are larger than Δ*R*.

#### 4.2.5 Describing full syntactic dependencies with PSTs

To describe phrase sequence syntactic dependencies of orders higher than 2nd we use Probabilistic suffix trees (PSTs).

##### Model fitting

This method was described in Markowitz et. al. 2013, code in https://github.com/jmarkow/pst). Briefly, the tree is a directed graph in which each phrase type is a root node representing the first order (Markov) transition probabilities to downstream phrases, including the end of song. The pie charts in Fig. 4C show such probabilities. Upstream nodes represent higher order Markov chains that are added sequentially if they significantly add information about the transition.

##### Model cross validation

To prevent overfitting, nodes in the probabilistic suffix trees are added only if they appear more often than a threshold frequency, *P*_*min*_. To determine *P*_*min*_ we replicate the procedure in [31] and carry a 10-fold model cross validation procedure. In this procedure the dataset is randomly divided into a training set, containing 90 percent of songs, and a test set, containing 10 percent of songs. A PST is created using the training set and used to calculate the negative log likelihood of the test set. This procedure is repeated 10 times for each value of *P*_*min*_. For data sets of different sizes the mean negative log-likelihood across the 10 cross validation subsets and across 10 data sets is then used to find the optimal value of *P*_*min*_ - the minimum negative log-likelihood that corresponds to the highest precision without over-fitting the training set. All PSTs in this work are created using a cross-validated *P*_*min*_.

##### Estimating PSTs in batches of days

To describe changes in syntactic dependencies we carried a ’rolling regression’ analysis of PSTs. For each canary that was recorded for 11 days or more, we divided the recordings time to overlapping batches of 5 days. We chose a time step of 1 day so consecutive batches overlapped in 4 out of 5 days. We repeated the PST model fitting and cross validation in each batch. This process yielded a different threshold frequency, *P*_*min*_, for each batch. To avoid misleading change in detail due to the different *P*_*min*_s, we picked the maximal *P*_*min*_ across all batches and estimated the batch-wise PST using this *P*_*min*_ for all batches. For the 5 birds in this analysis, the *P*_*min*_ values were 0.004, 0.006, 0.008, 0.005, 0.004, 0.005. These values were also used for calculating a PST for the entire data of each bird for the purpose of estimating the number of history-dependent transitions for each bird.

##### Calculating the normalized number of PST nodes

The size of a bird’s repertoire of syllables, *N*_*syllables*_, directly determines how many branches can be in the PST that describe the bird’s syntactic dependencies. This number is exactly the number of potential 1^*st*^ order Markov transition probabilities (e.g. as shown in the inner circle of pies in 4E). Similarly, the potential number of 2^*nd*^ order dependencies scales like 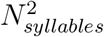 and, generally, the potential number of history dependencies of order *k* scales like 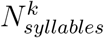

To compare the number of PST nodes across birds we accounted for the different syllable types produced by each bird by normalizing the number of k-deep dependencies, identified in the PST, by 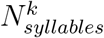 where *N*_*syllables*_ is the number of syllables produced by that bird. In 4C, we average this normalized number across birds and then display the fractional change from the first batch of days. Specifically, the fractional change of a quantity *x*(*n*) for *n* = 1..*m* is defined as 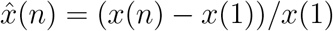

#### 4.2.6 Computing Ngram entropies

Phrase Ngrams are sequences of N consecutive phrases, also referred to as sequences of the N^th^ order. Not all possible Ngrams appear in the song of a canary and those that appear don’t have the same appearance likelihood. Using the same rolling 5-day data splits, described above, we estimate the probability distribution functions for Ngrams of orders 2-7 by directly measuring the frequency of each sequence. For example, the frequency of the 3-gram ’ABC’ is the number of times ‘ABC’ appears in the dataset divided by the total number of 3-gram appearances.

We calculate the Shannon entropy for each Ngram probability distribution function, 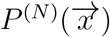 using 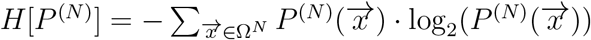 where Ω is the set of phrase types.

These entropy values are reported, for example, in 4B. Since birds have different numbers of phrase types, we average across birds after subtracting each bird’s entropy of 2-grams from the other Ngram orders before taking the average (4A).

---

**Supplementary Figure 1:**
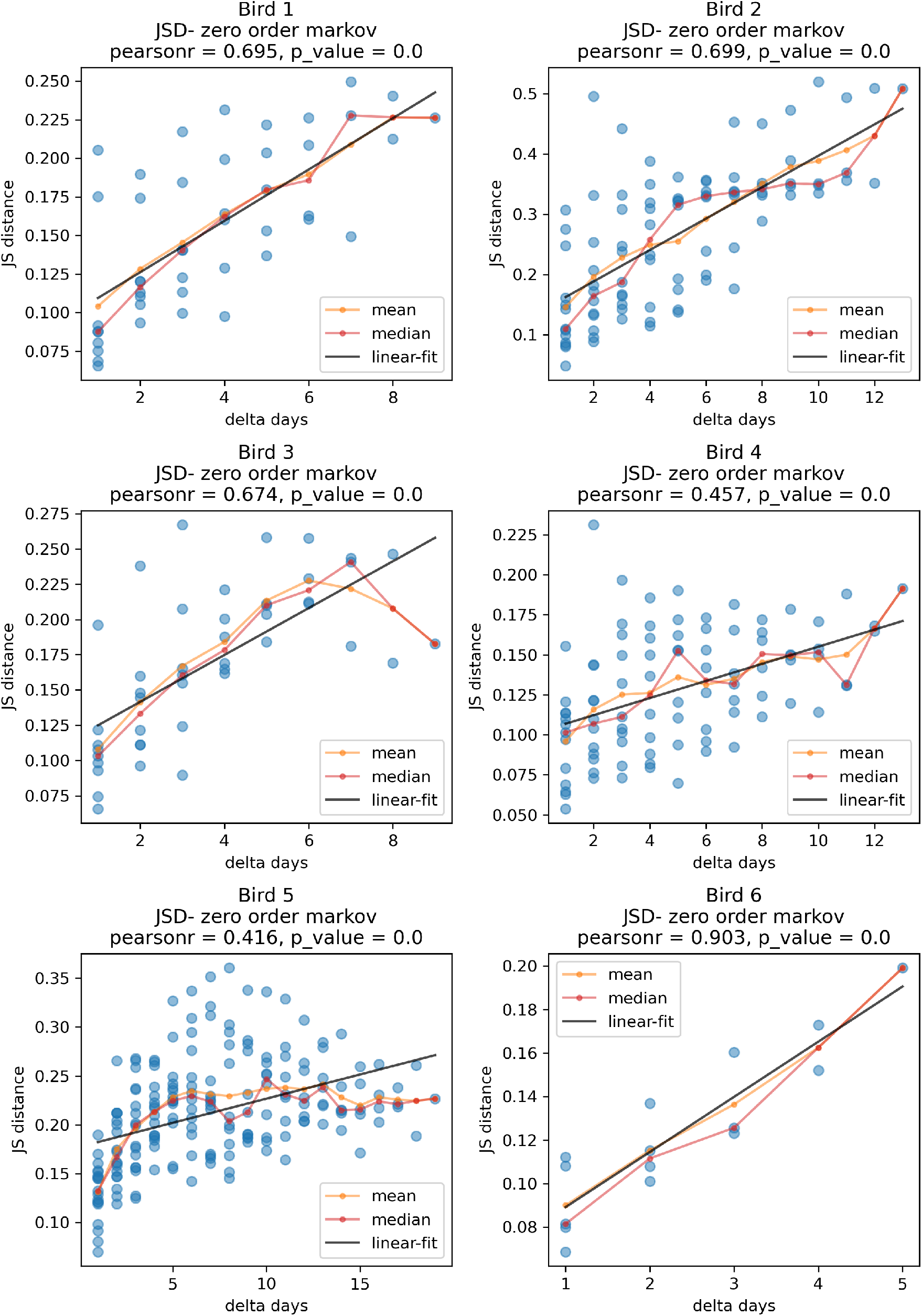
Phrase type use distributions move farther apart across days. Panels show results for 6 birds. Dots show the distance 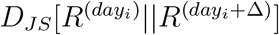 as a function of Δ. Lines show the linear trend (black) and the running mean and median (orange, red). Pearson’s r,p test for significant correlations.

**Supplementary Figure 2:**
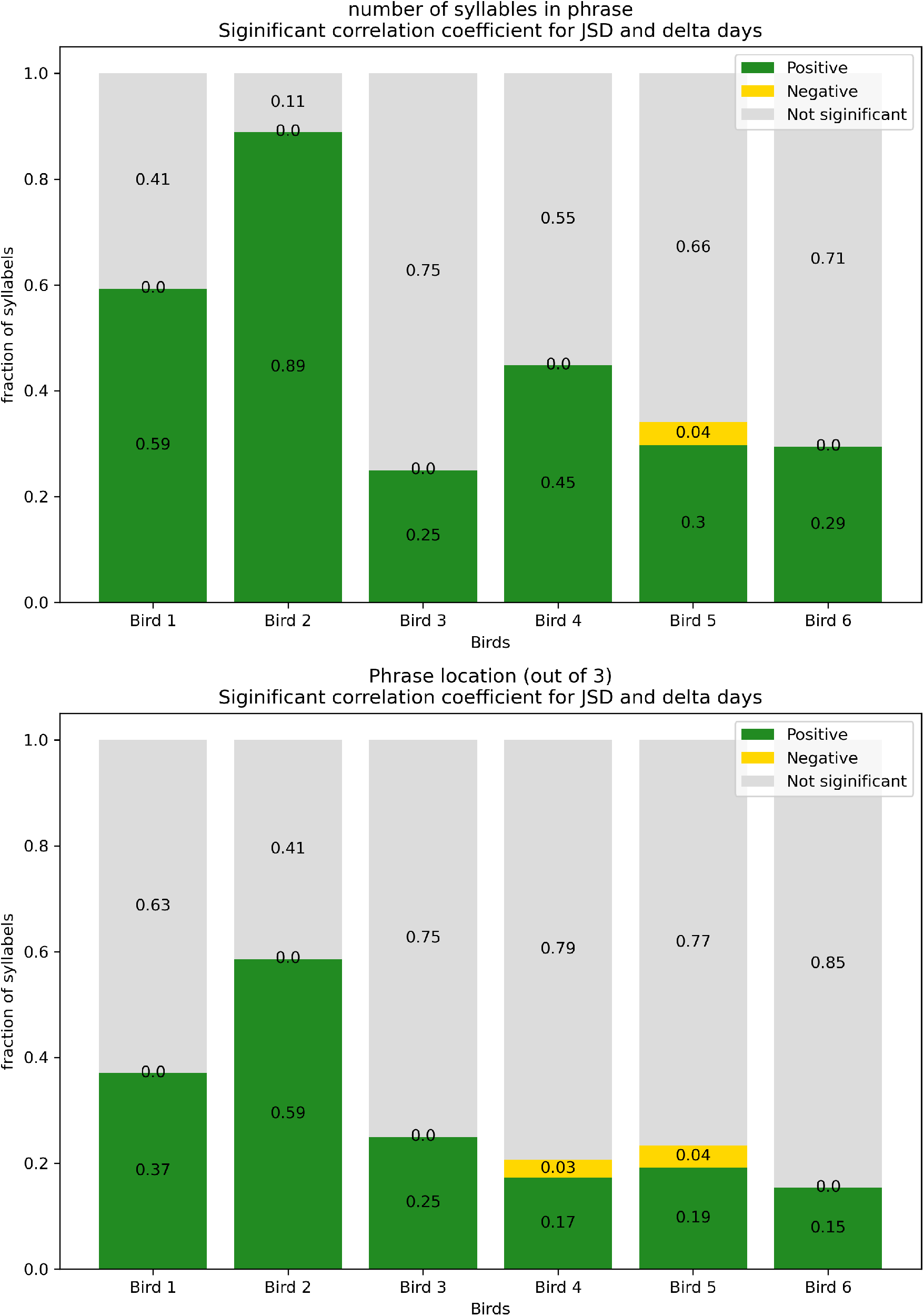
Fraction of distributions of syllable repeat numbers in phrases and phrase position in song that move apart across days. Green and yellow bars show fractions of positive and negative significant Pearson correlations (*p <* 0.025). **Top**, correlations between 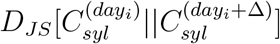 and Δ. **Bottom**, correlations between 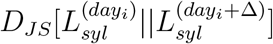 and Δ

**Supplementary Figure 3:**
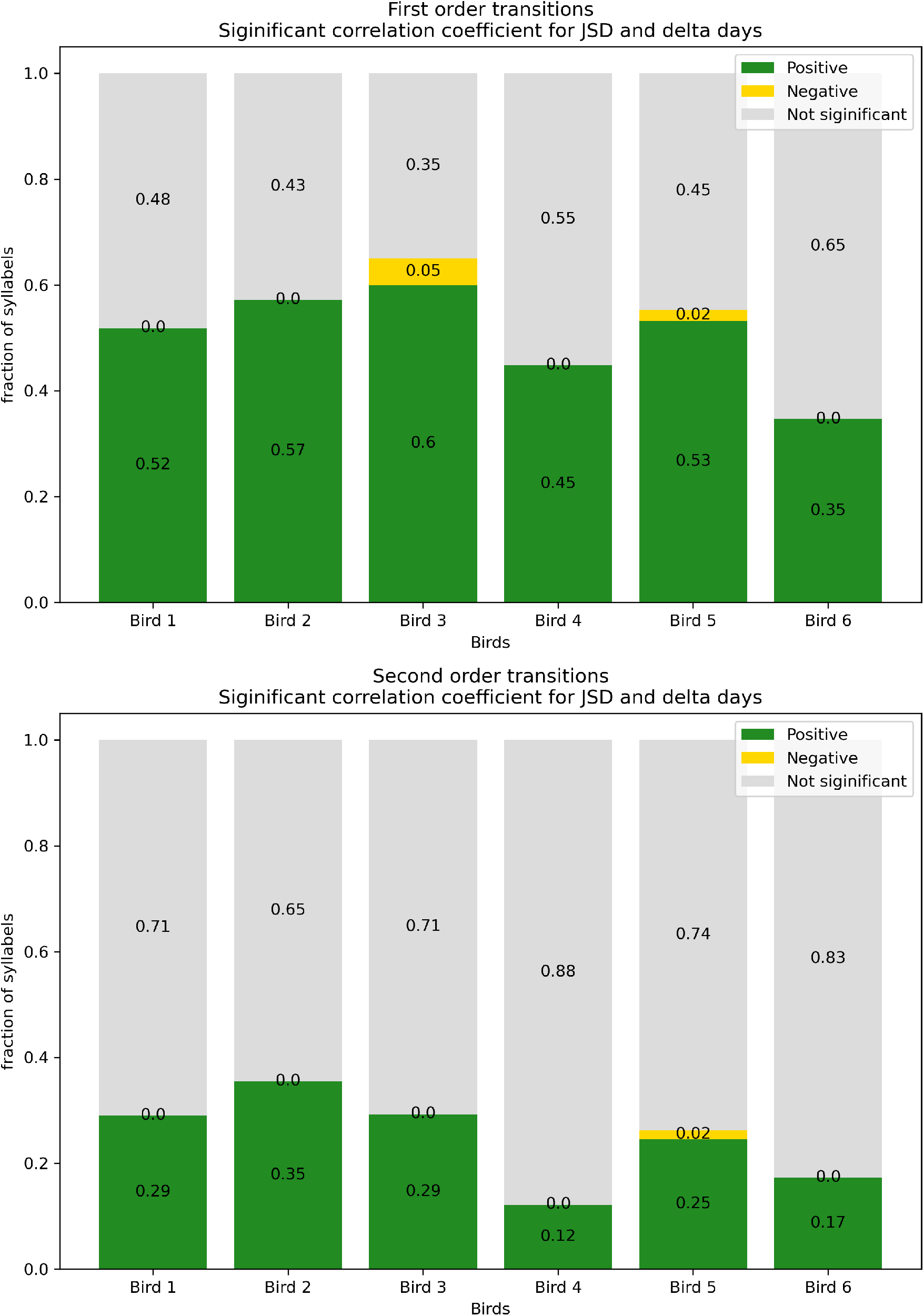
Fraction of phrase transitions that move apart across days. Green and yellow bars show fractions of positive and negative significant Pearson correlations (*p <* 0.025). **Top**, correlations between 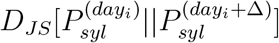 and Δ. **Bottom**, correlations between 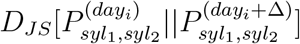 and Δ

**Supplementary Figure 4:**
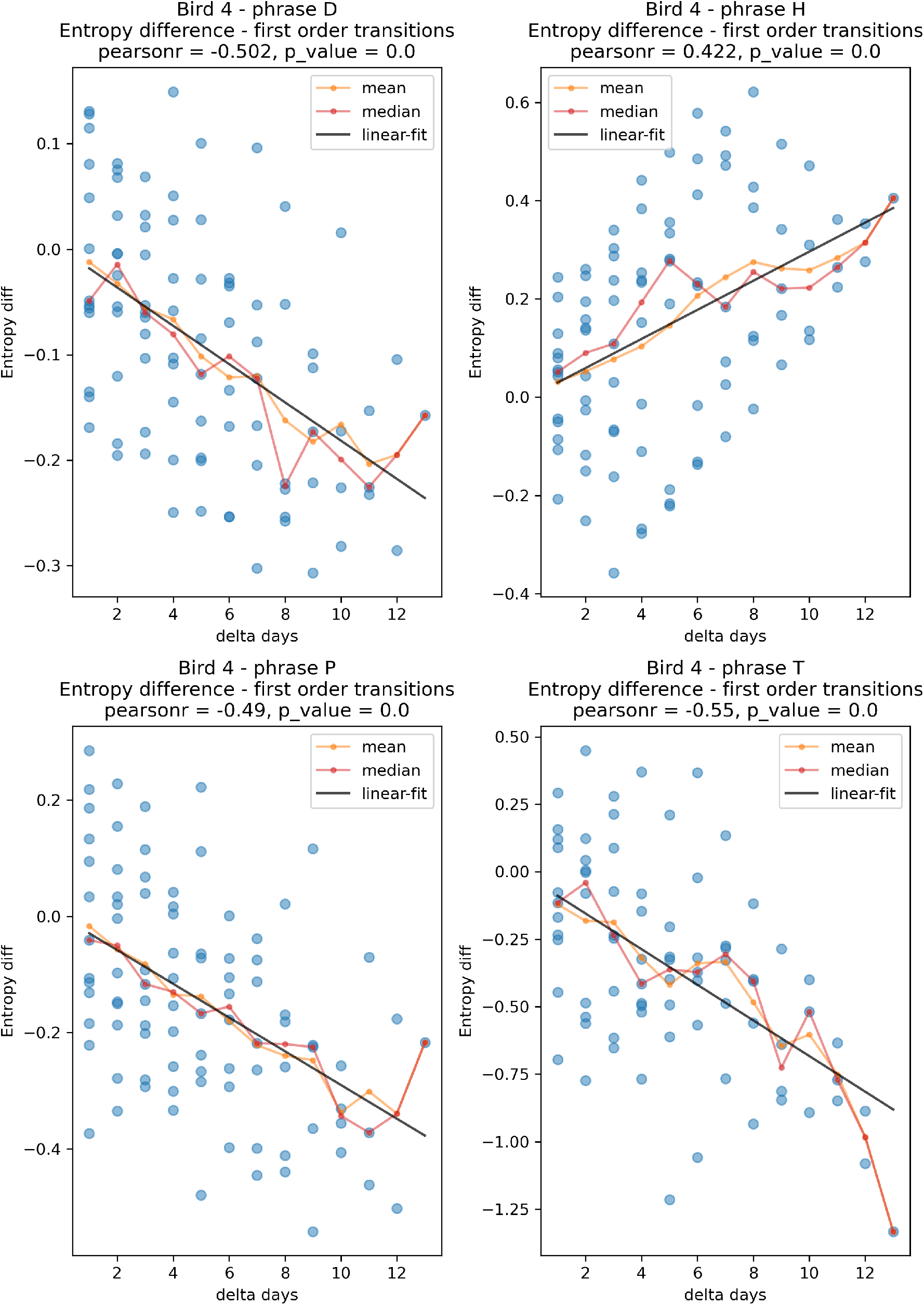
Example 1^st^ order transitions that change in entropy in correlation to the difference in days. Dots show the entropy difference, 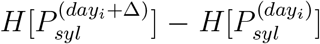 as a function of Δ. Lines show the linear trend (black) and the running mean and median (orange, red). Pearson’s r,p test for significant correlations.

**Supplementary Figure 5:**
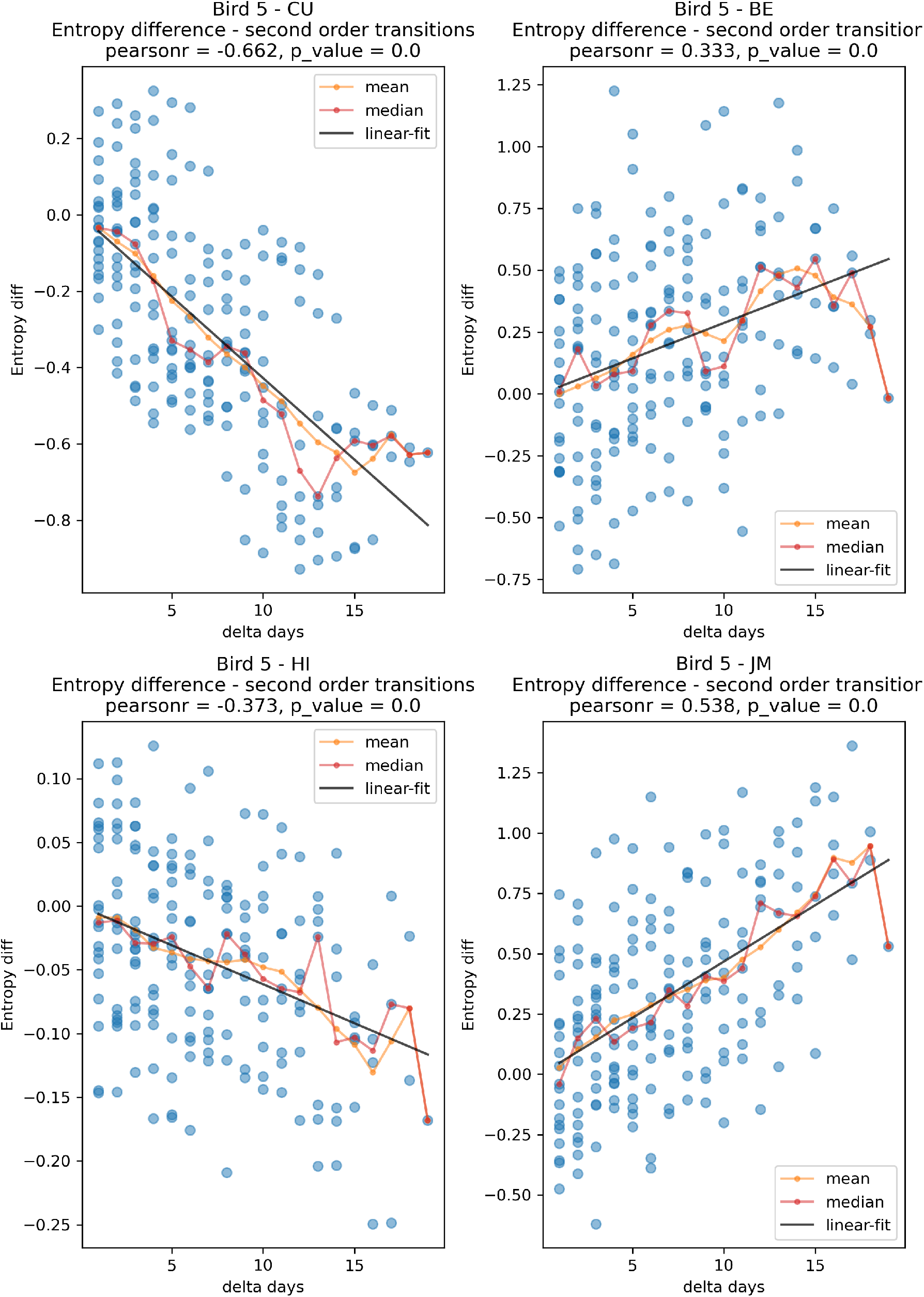
Example 2^nd^ order transitions that change in entropy in correlation to the difference in days. Dots show the entropy difference, 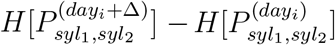 as a function of Δ. Lines show the linear trend (black) and the running mean and median (orange, red). Pearson’s r,p test for significant correlations.

**Supplementary Figure 6:**
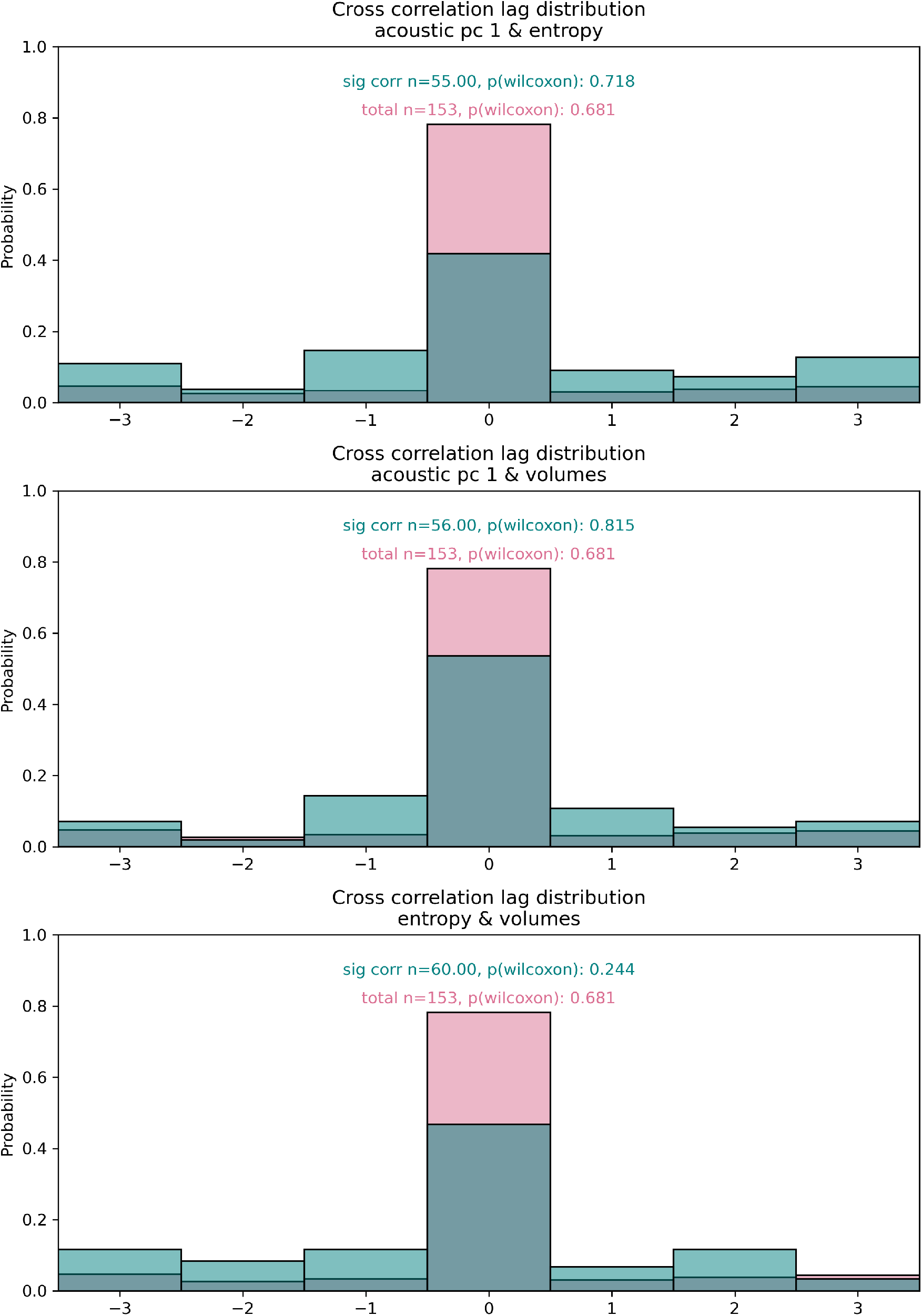
Lags between acoustics and 1^st^ order transitions. Panels show the distributions of optimal lag estimations (using cross correlations) between 3 variables: The entropy of 1^st^ order transitions from phrase type ’syl’ 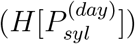, the average first principal coordinate in acoustics space of the syllable 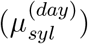 and the syllab’es acoustic variability 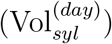 Blue histograms show values for only significant Pearson correlations (*p <* 0.05) at the optimal lag. Red histograms show values for all optimal lags. n values are for all 6 birds. Wilcoxon’s signed rank tests are used to evaluate the null hypotheses of no lag (p-values, all larger than 0.2).

